# BMI-CNV: A Bayesian framework for multiple genotyping platforms detection of copy number variation

**DOI:** 10.1101/2021.06.22.449433

**Authors:** Xizhi Luo, Guoshuai Cai, Alexander C. Mclain, Christopher I. Amos, Bo Cai, Feifei Xiao

## Abstract

Whole-exome sequencing (WES) enables detection of Copy number variations (CNVs) with high resolution in protein-coding regions. However, variations in the intergenic or intragenic regions are excluded from studies. Fortunately, samples have been previously sequenced by other genotyping platforms, such as SNP array. Moreover, conventional single sample-based methods suffer from high false discovery rate due to prominent data noise. Therefore, methods for integrating multiple genotyping platforms and samples are highly demanded for improved CNV detection.

We developed BMI-CNV, a <u>B</u>ayesian <u>M</u>ulti-sample and <u>I</u>ntegrative CNV (BMI-CNV) profiling method with data sequenced by both WES and microarray. For the multi-sample integration, we identify the shared CNVs regions across samples using a Bayesian probit stick-breaking process model coupled with a Gaussian Mixture model estimation. With extensive simulations, BMI-CNV outperformed existing methods with remarkably improved accuracy. By applying to the matched 1000 genomes project and HapMap project data, we showed that BMI-CNV accurately detected common variants. We further applied it to The Research of International Cancer of Lung (TRICL) consortium with matched WES and OncoArray data and identified lung cancer risk associated genes in 17q11.2, 1p36.12, 8q23.1 and 5q22.2 regions, which may provide new insights into the etiology of lung cancer.

## 1. Introduction

Copy number variations (CNVs) refers to the phenomenon that the copy number of specific genomic segments varies among individuals. CNVs are a major type of structural variation comprised of deletions and duplications on the genomic segments. They have been found to play an important role in complex diseases such as cancer, muscle diseases, and neuropsychiatric diseases (1–3). Also, a recurrent deletion at 17q12 has been revealed to be associated with an increased risk of autism spectrum disorders and schizophrenia (4). A common CNV in the *Hp* gene has been discovered to be a risk factor for cardiovascular disease in patients with type 2 diabetes (5).

With the dramatic growth of modern technologies and the accompanying cost drop in sequencing, massive whole-exome sequencing (WES) datasets have been generated from large-scale biomedical studies, which allows for the identification of genomic variants in functional protein-coding regions (6). However, exons only encompass 1% of the genome, limiting the possibility to investigate the impact of CNVs located in the non-coding regions (7). Moreover, WES is subject to the non-uniform coverage of sequence reads in the assembly procedure due to the existence of short duplications or deletions, resulting in many dropped out segments which are originally mapped to the exome. For example, Fang et al., found that more than 16% of the exons cannot be captured by WES experiments (8). This will lead to negligence in detecting short CNVs and therefore integrating available SNP array has great potential to overcome this challenge. In many of these WES cohorts, the same samples have been previously genotyped by the SNP array. For instance, the international Transdisciplinary Research In Cancer of the Lung (TRICL) consortium genotyped 2,003 subjects with both WES and SNP array data (6). The Alzheimer’s Disease Genetics Consortium (ADGC) and Alzheimer’s Disease Sequencing Project (ADSP) (9, 10) have also collected such multi-platform data. Consequently, the demand for multi-platform (e.g., WES and SNP) integration methods, which will comprehensively study CNV in a full-coverage manner, has dramatically increased.

Similar efforts such as iCNV have been made by Zhou et al. for integrative segmentation (11). In iCNV, data from different platforms were first normalized and standardized and then jointly segmented by a Hidden Markov Model (11). This method had a significant boost in accuracy compared to WES, however, it only used information from a single sample. As we know, technological and biological factors prominent in real data usually increase the variations and noise in data intensities, leading to unreliable findings with single-sample scanning of CNVs. Consequently, multiple sample strategies previously introduced for detecting common CNVs can improve the robustness and detection power with noisy data. Such a direction has been supported by various existing studies (12–14), but none has focused on multi-platform integration. Moreover, most widely used WES methods such as CODEX2 and EXCAVATOR are also for single-sample scanning (15, 16). As a result, it is of highly demand to develop a full spectrum CNV detection method that can meanwhile achieve high accuracy using comprehensive information from all samples.

In this study, we developed BMI-CNV, a <u>B</u>ayesian <u>M</u>ulti-sample and <u>I</u>ntegrative CNV calling method. Comparing to existing approaches, BMI-CNV efficiently integrates data from multiple platforms (i.e., WES and SNP array) and multiple samples to exploit the comprehensive information, leading to a full spectrum study of CNVs. Extensive numerical simulation studies showed that BMI-CNV presented significantly improved sensitivity over the existing methods. The method was further illustrated by applying to the HapMap project (17) and the 1000 Genomes project (18). It was further applied to the international TRICL dataset to identify lung cancer-associated CNVs. This new method has a wide scope of applications and has great potential to be further extended to profile CNVs for whole-genome sequencing and single-cell sequencing data analyses.

## 2. Method

Our method mainly focuses on CNV detection by integrating the SNP array and WES data, although it can also be naturally applied to the WES data only situation. Figure 1 shows an overview of the framework of BMI-CNV. First, WES read counts and SNP array intensities are integrated using a series of data integration procedures, including normalization, standardization, and merging. Our main algorithm consists of two main stages: Stage I uses a Bayesian PSBP method (Section 2.1) coupled with a Gaussian mixture model-based initial data filtering (Section 2.3) to identify shared CNV regions, and Stage II as the individual CNV calling procedure (Section 2.2).

**Figure 1.**
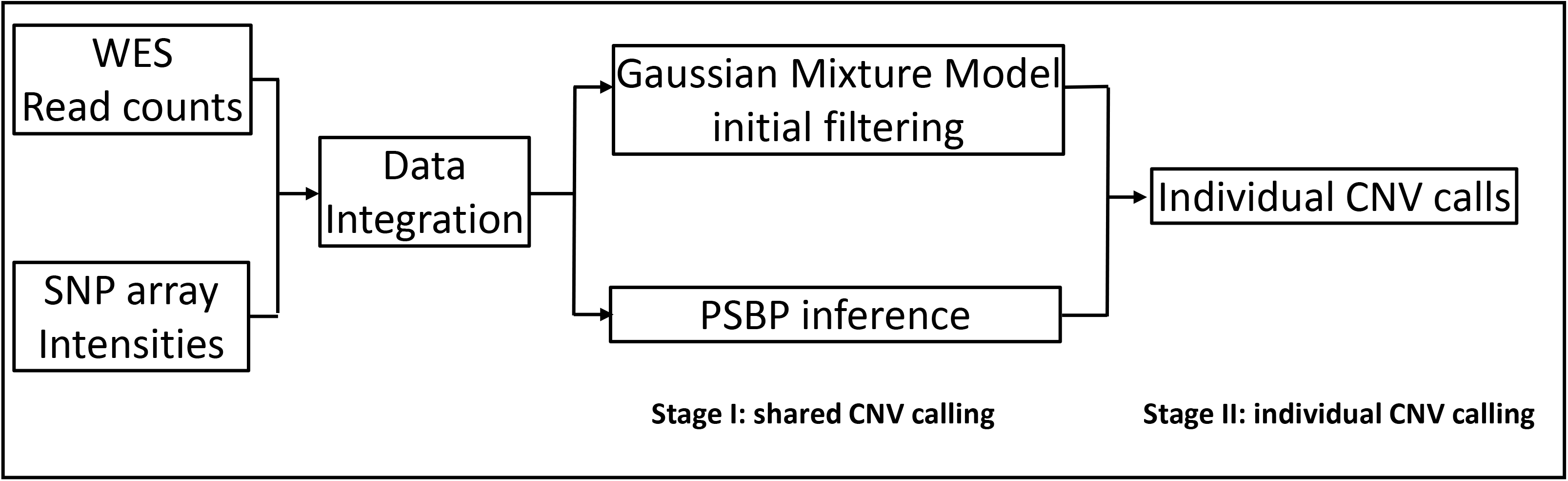
Analysis workflow of BMI-CNV. BMI-CNV requires two inputs: (1) WES raw read count data from testing and control samples that are computed by using genotyping tools such as SAMtools; (2) SNP array intensities. WES read counts are normalized to correct exon length, GC-content and mappability biases. Logarithm of normalized values between testing and pooled control samples are calculated. SNP array intensities are normalized to adjust the genomic waves. The WES and SNP array data are standardized by robust scaling approach and then integrated. For CNV calling, BMI-CNV carries out a two-stage framework to generate CNV calls. In stage I, an initial data filtering procedure is coupled with a Bayesian PSBP method to identify shared CNV regions. In stage II, an individual CNV calling procedure is performed to call CNVs in each sample.

### Data description, notations, and models

First, we performed platform-specific normalization procedures for the original data. For WES data, with raw read depth data from test and control samples, we adopted the exon mean read count (EMRC) introduced by Magi et al. as our input (15). We utilized the EXCAVATOR2 median normalization procedure, with external controls being pre-specified by researchers (19). The median normalization includes three steps to mitigate the effects from three observed sources of bias: exon length, GC-content, and mappability. We pooled all control samples by averaging the EMRC on each exon across all samples and calculated the logarithm of normalized EMRC between the test and pooled control samples, referred to as the *log*_2_*R* intensities. This *log*_2_*R* data was then processed by the lowess-scatter plot procedure to adjust read-depth differences between testing and control samples and remove coverage dependent bias. For array data, PennCNV was used to adjust the genomic wave on genetic intensities (i.e., Log R ratio (LRR)) (20).

To bring the SNP array LRR values and the WES-derived *log*_2_*R* to the same scale, we standardized each data via a robust scaling approach (21). Compared to the conventional standardization method, the robust scaling approach used median and interquartile ranges to mitigate the influence from potential outliers and signals from double deletions (Details in Supplementary A.1). The WES and SNP array data were merged by chromosomal coordinates to effectively integrate information for joint segmentation.

Let Y denote a *n* × *m* data matrix obtained from the pre-CNV calling procedures described above, where *Y*_*ij*_ represented the processed genetic intensities (e.g., LRR from array or *log*_2_*R* from WES) for the *i-th* (*i*=1, 2,…, m) marker (e.g., SNPs from array or exons from WES) in sample *j* (j=1, 2,…, n). We assumed a classic normal kernel for *Y*_*ij*_,

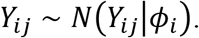

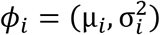 is an unknown vector of the underlying mean and variance at position *i* across all samples, in which different values of *ϕ*_*i*_ indicated the existence of different copy number states. We assumed there were five copy number states, including the deletion of a single copy (del.S), deletion of double copies (del.D), diploid, duplication of a single copy (dup.S), and duplication of double copies (dup.D). We considered τ to be a change point for sample *j* if *ϕ*_*j*,τ_ ≠ *ϕ*_*j*,τ+1_. The research goal is to estimate the locations of all the change points from all samples. CNV segments can then be generated by connecting adjacent change points. Conventional single sample methods have worked on this problem by simply applying the calling algorithm to each data sequence repeatedly (16, 19).

In our method, we assumed that certain change points were shared by multiple samples with population frequency *p*_*τ*_; and there were *G* change points in total. Let ***τ*** = {*τ*_1_, …, *τ*_G_} denote the locations of those shared change points. For each *τ*_*g*_(*g* = 1, … *G*), we considered the *j*-th sample as a CNV carrier if 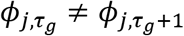. Therefore, the goal is now to estimate all the sample shared CNV regions (i.e., ***τ***) and then identify individual carriers. We used a two-stage method described below. Stage I uses a probit stick-breaking process to identify shared CNV regions (Section 2.1) and initially filters locations without CNVs (Section 2.3). Stage II calls CNVs individually (Section 2.2).

#### 2.1. Stage-I: shared CNV inference by Bayesian probit stick-breaking process model

We modelled the corresponding latent means and variances *ϕ*_*i*_ using the Bayesian model through a probit stick-breaking process (PSBP) (22, 23). The PSBP has a nice shrinkage property, allowing for efficiently clustering high dimensional *ϕ*_*i*_ to a small number of clusters (i.e., copy number states). Moreover, the PSBP mixture model can capture multimodal and heavy-tailed distribution, which relaxed the normality assumption of latent *ϕ*_*i*_, providing more flexible scenarios for modelling the complex CNV data. Specifically, we assumed *ϕ*_*i*_ followed an unknown distribution *G* ∼ *PSBP*(α*G*_0_) with centering distribution *G*_0_ and shape measure *α* that reflected how far away the random distribution *G* is from the center *G*_0_. Following Rodriguez et al., *G* admitted a representation of the form (23):

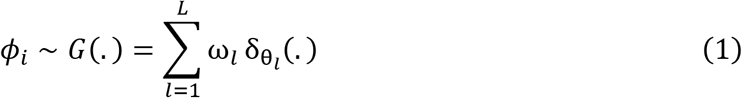

where *L* represented the number of all possible copy number states (e.g., L=5), 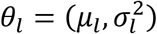 are possible distinct mean and variance specific to each copy number state (*l* = 1, 2, ‥, *L*), 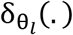 is a degenerate distribution at *θ*_*l*_, and *ω*_*l*_ = Φ(α_*l*_) ∏_*r*<*l*_(1 − Φ(α_*r*_)) represented the probability of assigning *θ*_*l*_ to each position where Φ(.) is the probit function and 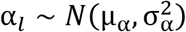. Following this structure, each *ϕ*_*i*_ was assigned to one of the {*θ*_*l*_} based on the observed intensities across all potential carriers of the copy number state for locus *i*. The carriers were initially identified using the strategy described below in Section 2.3. To simultaneously implement the variable selection and clustering procedures for the purpose of CNV detection, we further reconstructed the PSBP model as follows (24, 25):

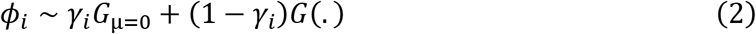

where the *G*_μ=0_ was the underlying distribution of the normal copy number states with the mean fixed at zero (i.e., diploids). *γ*_*i*_ ∼ *Bernoulli*(κ) is an indicator of *ϕ*_*i*_ being in *G*_μ=0_ (i.e., normal state) or not, which incorporated variable selection of the locus across samples. Specifically, when *γ*_*i*_ = 1, *ϕ*_*i*_ followed a distribution *G*_μ=0_; whereas *γ*_*i*_ = 0 indicated a potential CNV locus following *G*(.) defined in equation (1). Within this framework, the posterior probability of *ϕ*_*i*_ being *G*_μ=0_ or not was first calculated through inference on *γ*_*i*_. While the *ϕ*_*i*_ given *γ*_*i*_ = 0 was then assigned to its most possible state according to the posterior probabilities of belonging to various copy number states (details in Section 2.4).

#### 2.2. Stage-II: Individual CNV calling

After a CNV region is identified across samples, we then determine the carriers in the samples, that is, to call CNVs in each individual sample. Specifically, after obtaining the posterior mean and variance estimates specific to each copy number state (i.e., 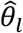 and 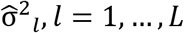), we constructed the interval for each state as 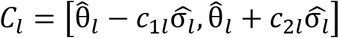. We considered a sample as the carrier and classified the segment into the *l-*th copy number state if its segmental mean fell within one specific interval *C*_*l*_. Values of *c*_1_ and *c*_2_ could be arbitrarily chosen according to empirical evidence about the magnitude of mean shifts of each CNV state, which may vary by genotyping platforms. In practice, we will suggest plotting the genotyping signals of CNV segments that were identified under different combinations of *c*_1_ and *c*_2_ for visualization. True positive rate (TPR) could be calculated for each combination. The optimal choices for *c*_1_ and *c*_2_ would achieve the highest TPR.

#### 2.3. Initial data pooling by Gaussian mixture model

A major complexity of multi-sample integration is how to effectively combine the information in the presence of samples containing no CNV signals. The intensities from non-carriers will dilute the signal, which may hence significantly decrease the detection power. Li and Tseng adopted a weighted technique to downweigh the non-carriers; Sung et al. used the ordered p-values from all samples and only selected samples with small p-values for CNV inference (14, 26). With the same spirit, we considered a preliminary data filtering step so that only intensities that were most likely arising from the carriers would be selected for stage-I CNV calling.

Specifically, for the *i-th* position, we let *Y*_*ij*_ = (*Y*_*i*1_, *Y*_*i*2_, …, *Y*_*in*_)^*T*^ denote the CNV data vector across all samples. A Gaussian mixture model with three copy number mixture states (including deletions, normal state, and duplications) was considered:

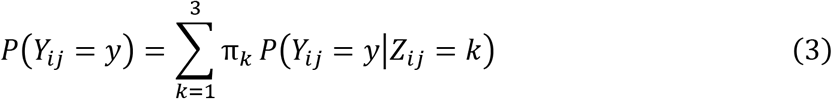

*Z*_*ij*_ was the latent mixture component, π_*k*_ was the mixture proportion reflecting the probability that *Y*_*ij*_ belonged to the *k*-th mixture component, and *P*(*Y*_*ij*_ = *y*|*Z*_*ij*_ = *k*) ∼ *N*(μ_*k*_, σ^2^_*k*_) was the component distribution. All the component-specific parameters (i.e., π_*k*_, μ_*k*_, σ^2^_*k*_) were estimated by the expectation-maximization (EM) algorithm. Samples were then assigned to latent clusters with the largest estimates of π_*k*_ (27). Afterward, samples showing evidence of diploid (i.e., *k* = 2) will be filtered.

#### 2.4. Hyperparameters choices and MCMC algorithm

For the PSBP mixture model described in equations (1–2), we developed a Markov Chain Monte Carlo (MCMC) algorithm relying on a modification of the Gibbs sampler to perform the posterior inference (28). Note that with the proper choices of priors and hyperpriors described below, all full conditionals are very straightforward and can be analytically derived.

We adopted the following choices for the hyperparameters (Details in Supplementary A.2). For variable selection variable *γ*_*i*_ ∼ *Ber*(κ), we used the Beta conjugate hyperprior for the parameter κ. For the component-specific 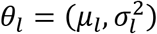 of *G*(.) defined in equation (1), we used the conjugate normal and inverse gamma hyperpriors for *μ*_*l*_ and 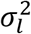, respectively. The posteriors of parameters can then be computed via the MCMC algorithm, for which the detailed updating steps are in Supplementary A.3. We introduced the latent variable *s*_*i*_ such that *s*_*i*_ = *l* denoted the *i*-th position was assigned to the *l-*th component, and *s*_*i*_ was sampled from a multinomial distribution. To update the latent α_*l*_ and weight parameter *ω*_*l*_, we adopted a data augmentation approach (23). We introduced a collection of conditionally independent latent variable *Z*_*il*_(*s*_*i*_) ∼ *N*(α_*l*_, 1), and we defined *s*_*i*_ = *l* if and only if *Z*_*il*_(*s*_*i*_) > 0 and *Z*_*ir*_(*s*_*i*_) < 0 for r < l. Therefore, we have,

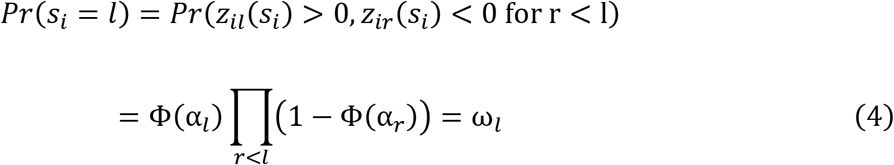

The augmented variable *Z*_*il*_(*s*_*i*_) can be imputed by sampling from its full conditional distribution,

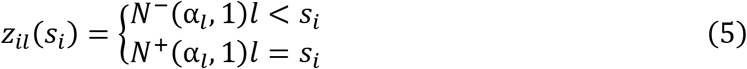

where *N*^−^ and *N*^+^ denoted the negative and positive truncated normal distributions. Given *Z*_*il*_(*s*_*i*_), α_*l*_ can be updated from its conjugate full conditional distribution. The component-specific parameters, *θ*_*l*_ and σ^2^_*l*_ were also updated from their conjugate full conditionals. Finally, we updated *γ*_*i*_ based on the marginal likelihoods for (***y***, ***s***) and *κ* from its conjugate full conditional distribution.

#### 2.5. Numerical Simulations

To evaluate the performance of our method, we conducted simulations under various settings. Four copy number states were simulated including del.S, del.D, dup.S, and dup.D. The CNV length varied from 10~30 markers (i.e., SNPs and exons), 30~60 markers and 60~100 markers. The CNV population frequency was set to be 20%, 50% or 100%, respectively.

First, we evaluated our method when both WES and SNP array data were available. For WES data, to generate data retaining the true noise structure and exon distribution, we conducted a spike-in design (11, 16). We started with read depth data on chromosome 1 in 81 samples from the 1000 Genomes Project (18). Exons harboring CNVs identified by EXCAVATOR2 and CODEX2 and reported in the Database of Genomic Variants (DGV) were removed (16, 19, 29). The read depth data of the remaining exons were treated as WES random noise background. We multiplied the background read depth by *c*/2, where *c* was sampled from a normal distribution with mean and variance provided in Supplementary A.6. For SNP array data, we utilized the similar strategy used in Xiao et al. to simulate intensities (i.e., LRR) (30), which mimicked the real data from the international HapMap consortium (17). We then randomly selected and spiked in 50 dispersed CNV segments of varying length and frequency for every single sequence.

Using the simulated datasets, our method BMI-CNV was compared to the existing integrative method iCNV (11). The performance of methods was assessed by precision rate, recall rate, and F1 score measures (Supplementary A.6). We also evaluated the performance of our method when only WES data was available. Our method’s performance was compared against that of CODEX2, EXCAVATOR2, and iCNV in the single-platform mode (11, 16, 19).

#### 2.6. Application to the 1000 Genomes project and HapMap datasets

To further illustrate the characteristics of our method, we analyzed the same 81 individuals using SNP array and WES data from the 1000 genomes project and the international HapMap consortium. A detailed description of the experimental samples and genotyping platforms was provided in previous literature (17, 18). Raw read counts and SNP array data were processed and normalized to generate *log*_2_*R* and LRR intensities. For the WES data, we arbitrarily selected four normal samples from the 1000 genomes project as controls in the calculation of *log*_2_*R* intensity (details in Supplementary A.4). The posterior inference of BMI-CNV was based on 2,000 MCMC samples with a burn-in period of 500 iterations.

#### 2.7. Integrative Analysis of TRICL case-control study

We further applied BMI-CNV to the international lung cancer study TRICL (6). We applied BMI-CNV to 1,163 samples genotyped by both OncoArray and WES data, and 829 samples that only had WES data using our integrative analysis mode and single platform analysis mode, respectively (details in Supplementary A.4 and A.5).

CNV calls were annotated by known gene regions obtained from UCSC Genome Browser (31). A gene-based association test was performed to investigate the influence of CNV on lung cancer susceptibility:

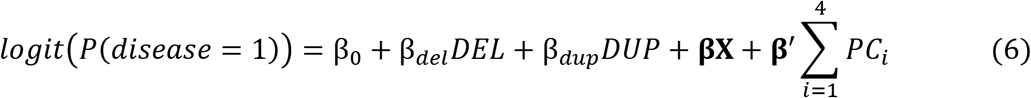

*DEL* and *DUP* were two indicator variables for deletions and duplications separately. We adjusted the covariates, including smoking status (ever/never), age, and gender. *PCs* were the top 4 principal components representing the ancestry information (32). In addition to study the overall lung cancer risk, we also performed stratification analyses by histological types of lung cancer (squamous cell lung cancer [SQC] and lung adenocarcinoma [LUAD]). The significance of the effects from deletions and duplications were tested separately via the Wald test (i.e., β_*del*_ = 0, β_*dup*_ = 0), all nominal P-values were adjusted by Benjamini-Hochberg (BH) procedure (33).

## 3. Results

### 3.1. Simulations showed improved performance of BMI-CNV in integrative analysis

We first evaluated the performance of BMI-CNV with simulated data when both SNP array and WES data were available. Various simulation settings were considered, including different CNV sizes and population frequencies. Overall, BMI-CNV outperformed iCNV in all scenarios with higher F1 scores (Figure 2, Table 1). iCNV tended to be conservative compared to our method, which maintained a high precision rate, although the recall rate was compromised. For example, when the simulated CNVs had a length of 30-60 markers and the population frequency was 20%, BMI-CNV had a precision at 0.70, a recall rate at 0.83, and an F1 score at 0.76. The corresponding values for iCNV were 0.99, 0.37, and 0.54, respectively. Moreover, at a certain value of CNV size, we observed the improved performance of BMI-CNV as CNV frequencies increased from 20% to 100%, and it achieved the highest F1 score when all the samples were carriers. The performance of the iCNV method was not sensitive to the CNV frequencies. Regarding computational speed, our method took about 280 minutes to scan a chromosome with 90,739 markers from 81 samples based on 2,000 MCMC sampling runs. The computation was performed on a regular laptop with an Intel Core i7 processor and 24.00 GB of RAM.

**Table 1.** Summary of performance of our method on simulated data in the integrative analysis. Precision rate, recall rate, and F1 score are summarized for CNV calls generated by BMI-CNV and iCNV in the integrative analysis. Frequency: CNV frequency.

|  |  | BMI-CNV |  |  | iCNV |  |  |
| --- | --- | --- | --- | --- | --- | --- | --- |
| CNV length (markers) | Frequency | Precision | Recall | F1 | Precision | Recall | F1 |
| 10~30 | 20% | 0.54 | 0.71 | 0.61 | 0.98 | 0.35 | 0.51 |
|  | 50% | 0.58 | 0.90 | 0.71 | 0.98 | 0.35 | 0.51 |
|  | 100% | 0.79 | 0.91 | 0.84 | 0.98 | 0.35 | 0.52 |
| 30~60 | 20% | 0.70 | 0.83 | 0.76 | 0.99 | 0.37 | 0.54 |
|  | 50% | 0.75 | 0.87 | 0.81 | 0.99 | 0.35 | 0.52 |
|  | 100% | 0.92 | 0.85 | 0.89 | 0.99 | 0.36 | 0.53 |
| 60~100 | 20% | 0.75 | 0.88 | 0.81 | 0.99 | 0.40 | 0.57 |
|  | 50% | 0.87 | 0.89 | 0.88 | 0.99 | 0.39 | 0.56 |
|  | 100% | 0.88 | 0.89 | 0.89 | 0.99 | 0.40 | 0.57 |

**Figure 2.**
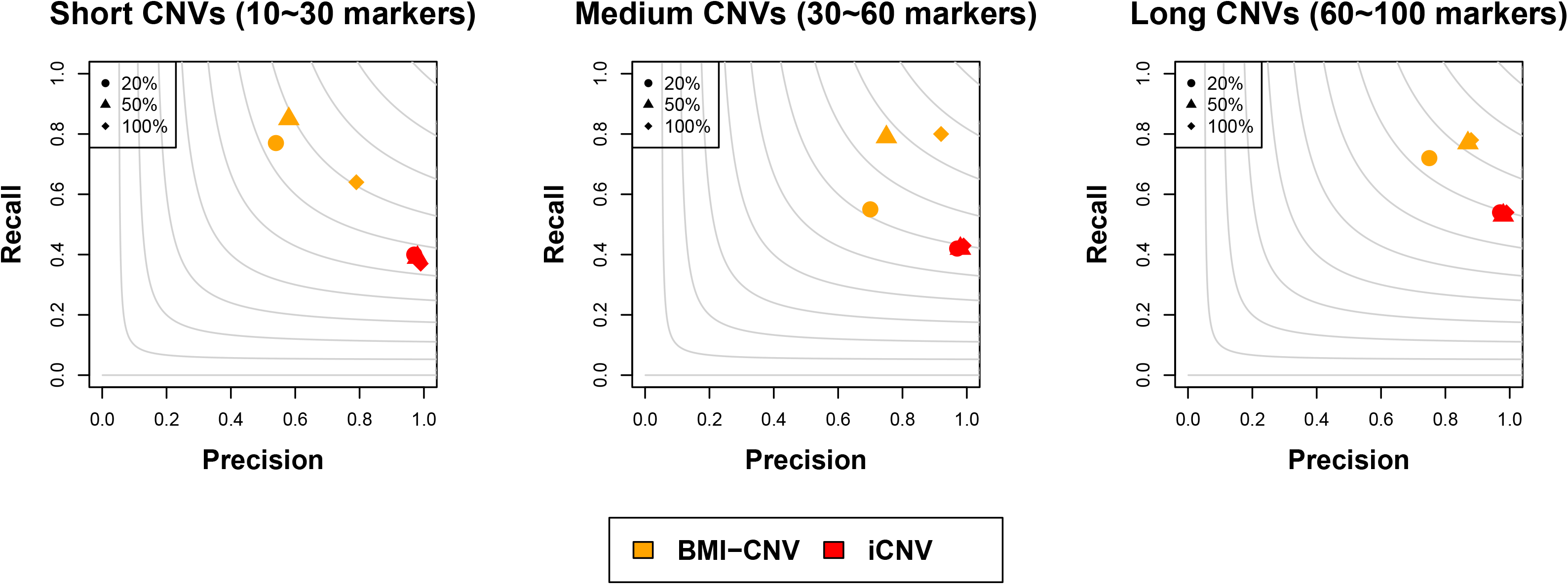
Performance assessment of BMI-CNV and iCNV on simulated data in the integrative analysis. Simulated CNVs are of frequency 20%, 50%, and 100% and length 10~30 markers (short), 30~60 markers (medium), and 60~100 markers (long). The grey contours are F1 scores calculated as the harmonic mean of precision and recall rates.

### 3.2. Simulations showed improved performance of BMI-CNV in single platform analysis

Next, we utilized the simulated WES data to assess the performance of BMI-CNV benchmarking againest existing WES CNV detection methods. The performance of BMI-CNV was superior to other methods on detecting medium and long CNVs reflected by the largest F1 scores (Figure 3, Table 2). CODEX2 performed the best for detecting short CNVs, except for detecting low-frequency CNVs (i.e., frequency=20%). iCNV and EXCAVATOR2 tended to be conservative, as they achieved a high precision rate but with a significant sacrifice on recall rate. It was also noteworthy to mention that, when the CNV size was fixed at a certain value, the performance of BMI-CNV and CODEX2 were both improved with increased CNV frequencies, and they achieved the highest F1 scores when all the samples were carriers. Still, the performance of EXCAVATOR2 and iCNV were not subject to CNV frequencies, as they mainly scan one sample at a time. The shared information from multiple samples was not utilized in the calling algorithms.

**Table 2.** Summary of performance of our method on simulated data in WES analysis. Precision rate, recall rate, and F1 score are summarized for CNV calls generated by BMI-CNV, iCNV, EXCAVATOR2, and CODEX2 in WES analysis. Frequency: CNV frequency.

|  |  | BMI-CNV |  |  | iCNV |  |  | EXCAVATOR2 |  |  | CODEX2 |  |  |
| --- | --- | --- | --- | --- | --- | --- | --- | --- | --- | --- | --- | --- | --- |
| CNV length (markers) | Frequency | Precision | Recall | F1 | Precision | Recall | F1 | Precision | Recall | F1 | Precision | Recall | F1 |
| 10~30 | 20% | 0.44 | 0.77 | 0.56 | 0.99 | 0.40 | 0.57 | 0.90 | 0.21 | 0.34 | 0.37 | 0.99 | 0.54 |
|  | 50% | 0.46 | 0.85 | 0.59 | 0.98 | 0.39 | 0.56 | 0.87 | 0.21 | 0.34 | 0.49 | 0.98 | 0.65 |
|  | 100% | 0.90 | 0.64 | 0.74 | 0.99 | 0.37 | 0.54 | 0.87 | 0.25 | 0.39 | 0.97 | 0.63 | 0.76 |
| 30~60 | 20% | 0.81 | 0.55 | 0.65 | 0.99 | 0.42 | 0.59 | 0.80 | 0.19 | 0.30 | 0.35 | 0.99 | 0.51 |
|  | 50% | 0.65 | 0.79 | 0.71 | 0.99 | 0.42 | 0.59 | 0.81 | 0.20 | 0.32 | 0.48 | 0.91 | 0.63 |
|  | 100% | 0.82 | 0.79 | 0.81 | 0.99 | 0.43 | 0.60 | 0.84 | 0.25 | 0.38 | 0.99 | 0.69 | 0.81 |
| 60~100 | 20% | 0.78 | 0.72 | 0.75 | 0.99 | 0.54 | 0.70 | 0.93 | 0.26 | 0.41 | 0.25 | 0.74 | 0.37 |
|  | 50% | 0.80 | 0.77 | 0.78 | 0.99 | 0.53 | 0.69 | 0.87 | 0.31 | 0.46 | 0.45 | 0.42 | 0.43 |
|  | 100% | 0.87 | 0.78 | 0.83 | 0.99 | 0.54 | 0.70 | 0.90 | 0.39 | 0.54 | 0.99 | 0.68 | 0.81 |

**Figure 3.**
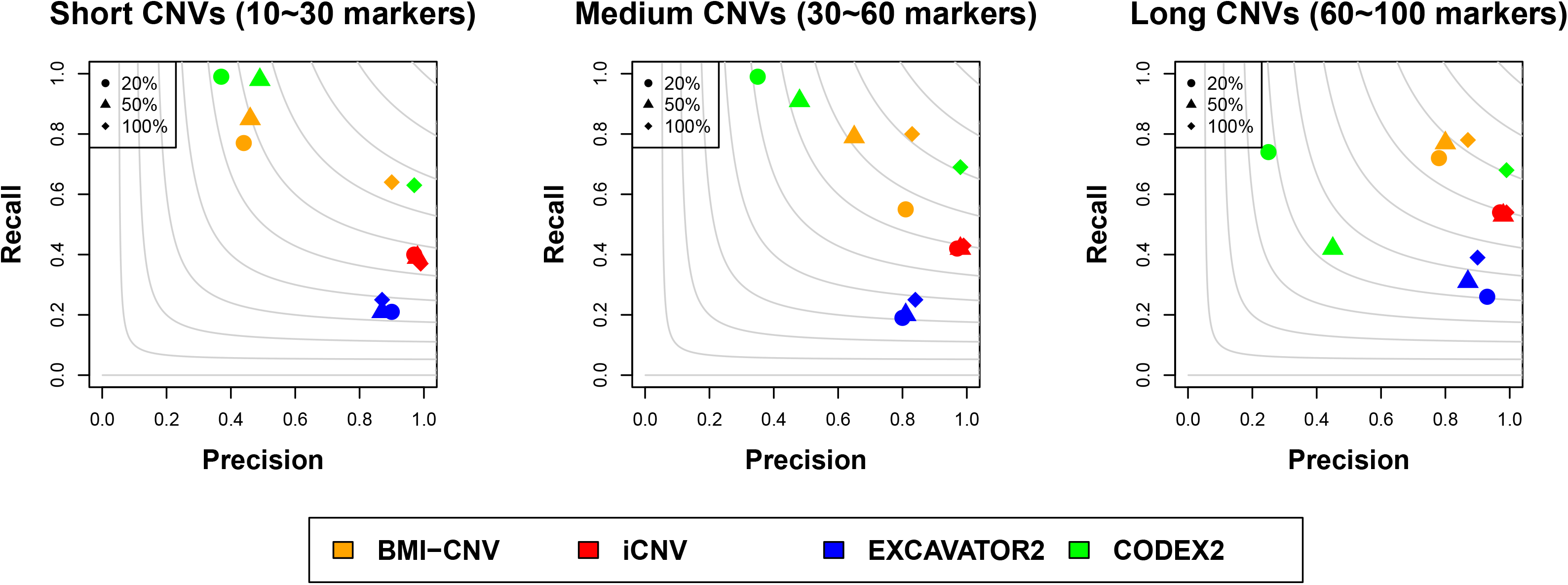
Performance assessment of BMI-CNV, iCNV, EXCAVATOR2, and CODEX2 on simulated data in WES analysis. Simulated CNVs are of frequency 20%, 50%, and 100% and length 10~30 markers (short), 30~60 markers (medium), and 60~100 markers (long). The grey contours are F1 scores calculated as the harmonic mean of precision and recall rates.

In conclusion, the BMI-CNV method, which integrated information from multiple samples, presented evidently improved performance in common CNV detection for both multi-platform integration and single-platform analyses.

#### Application to the 1000 genomes project and HapMap data

We further applied BMI-CNV to the public datasets from the 1000 genomes project and HapMap data for evaluation. In total, we identified 37, 213 CNVs from 81 samples (Figure 4). Among them, 28% of the CNVs have been previously reported by DGV (29). Most CNVs tended to be short (<20 markers) and had a frequency of less than 50%. The total number of deletions was nearly the same as that of duplications (19, 194 vs. 18, 019). Supplementary Figure S1 showed the summary of CNVs, which suggested no difference between deletions and duplications in the CNV length and frequency. Moreover, by integrating the SNP array data, our method recovered 4, 418 CNVs that resided in the non-coding regions that would be missed by the same method using WES data alone.

**Figure 4.**
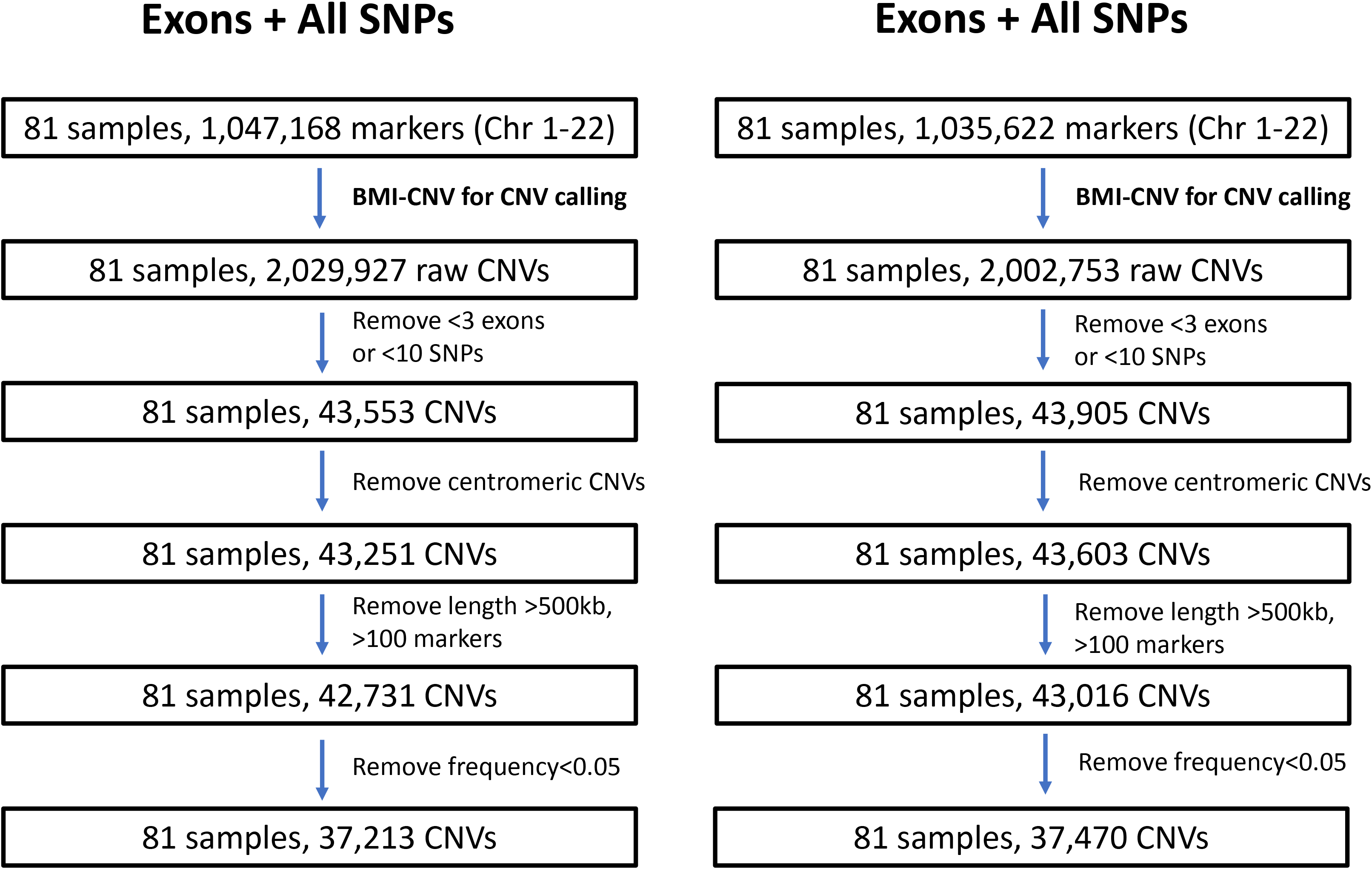
Overview of the application to the 1000 genomes project and HapMap data. The figure outlines the study design with a brief description of quality control (QC) methods. Summary of key results includes the sample size and number of CNVs at various stages of analysis. Left: CNV calling results using all SNPs and exons; right: CNV calling results using intronic SNPs and exons. Chr: chromosome.

Supplementary figure S2 illustrated one typical common deletion region, suggesting that 27 out of 81 samples were carriers of this variant. A clear pattern of the shared deletion was observed through visual inspection of the signal intensities for this region across samples, and our proposed method presented accuracy in identifying this region. We also explored an alternative data integration strategy that only used intronic SNPs from the array data. Comparing to the main method using all SNPs, integrative calling with only intronic SNPs yielded a similar number of CNVs but a lower concordance rate with DGV (26% vs. 28%), implying the advantage of utilizing information of all SNPs on slightly improved accuracy (Figure 4, Supplementary Figure S3).

### 3.3. Integrative Analysis of TRICL cases and controls

With the TRICL datasets, we identified 253,183 CNVs in autosomes from 1,992 samples (Figure 5) in total. Overall, we detected more deletions than duplications, with an average length of deletions (in markers) larger than that of the duplications (13.46 markers vs. 8.72 markers) (Supplementary Table S1). No significant difference in the overall proportion of deletions and duplications between cases and controls was observed (49% vs. 51% for deletions and 52% vs. 48% for duplications, respectively). Also, there was no difference in the length of deletions and duplications between cases and controls (Supplementary Table S1).

**Figure 5.**
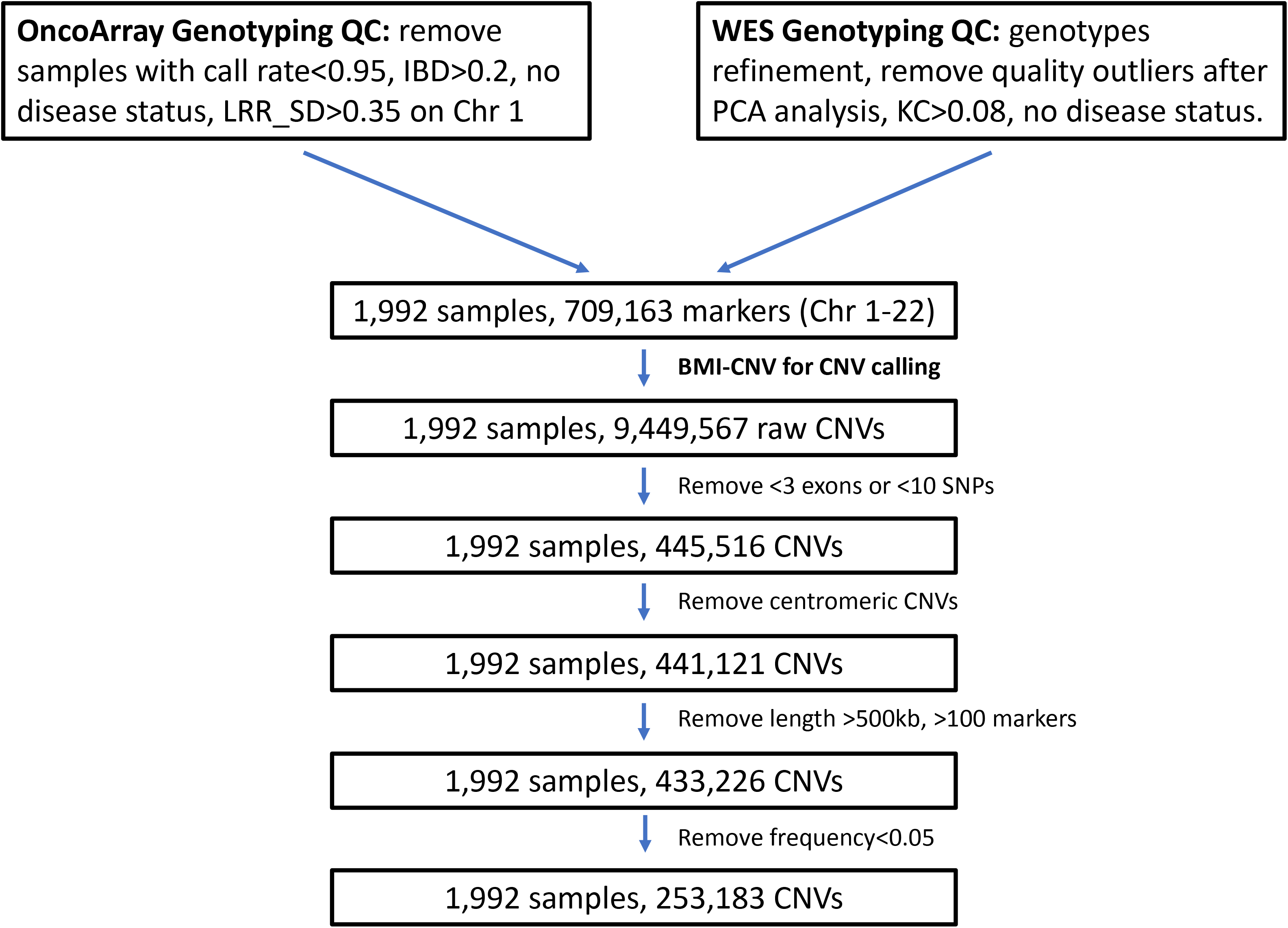
Overview of the integrative analysis of the TRICL case-control study. The figure outlines the study design with a brief description of quality control (QC) steps. Summary of key results includes the sample size and number of CNVs at various stages of analysis. IBD: identical by descent; KC: kinship coefficient; LRR_SD: standard deviation of Log R ratio; Chr: chromosome; PCA: principal component analysis.

Those identified CNVs were mapped to 3,472 genes. An association test with SQC subgroup highlighted the deletion gene *LGALS9* in 17q11.2 region (OR=4.14, 95% CI=1.65-10.38, *P*-value=0.002) and duplication genes *HSPG2* in 1p36.12 region (OR=4.79, 95% CI=1.75-13.10, *P*-value=0.002), *EIF3E* in 8q23.1 region (OR=2.19, 95% CI=1.31-3.64, *P*-value=0.003). Association results in the LUAD subgroup identified the duplication gene *YTHDC2* in 5q22.2 region (OR=2.88, 95% CI=1.62-5.12, *P*-value=0.0003), which was also identified in the overall lung cancer risk model by adjusting the histological subtypes as a covariate (Supplementary Table S2). The intensities’ plots indicated that all of those variants were valid CNV segments that showed distinct data patterns from other non-carriers and adjacent regions (Supplementary Figure S4). Although these genes were not significant after multiple comparison adjustments, they still provided potential evidence and great insights into future studies on the roles of CNVs in lung cancer risk.

## 4. Discussion

The importance of CNVs for elucidating the mechanism underlying many diseases has been increasingly remarked upon. Consequently, improving the accuracy of CNV detection is fundamental for downstream CNV-disease risk association and diagnostic classification. Nowadays, challenges shared by existing WES methods are the lack of sensitivity for common CNVs and the incapability of studying the non-coding regions of the genome. In this study, we have developed a multiple sample-based method, BMI-CNV, to improve common CNV detection with WES data, allowing for the integration of available SNP array data. The simulation results demonstrated the desirable performance across different scenarios of CNV sizes and population frequencies. The improvement for calling long and high-frequency CNVs was the most substantial. We reanalyzed the WES data from the 1000 Genomes Project and SNP array data previously generated by HapMap project 3 and demonstrated the advantage of multi-platform integration over the single-platform analysis. Finally, our application of BMI-CNV to WES and OncoArray datasets of the TRICL consortium indicated potential lung cancer associated CNVs.

This is the first report demonstrating the improved performance of CNV detection by utilizing a multi-sample and multiple-platform strategy. The advantages of our method in theory are in two aspects. First, utilizing information across samples will dramatically reduce the false positives and boost the detection power. We showed that BMI-CNV presented essential advantages over other single-sample methods in detecting common variants. The advantage has been previously shown in Song et al. (14), which proved that the underlying statistical power of multi-sample methods converged to one at a faster rate than single-sample methods. Second, BMI-CNV integrates available SNP data to detect CNVs in non-coding regions. allowing for full spectrum genomic variants investigation. Indeed, the important role of CNVs in non-coding regions has been revealed in numerous studies. For example, Kumaran et al. identified 1,812 breast cancer-associated CNVs mapping to non-coding regions (34). D’Aurizio et al. and Kuilman et al. developed WES-based methods, EXCAVATOR2 and CopywriteR, which used both the targeted reads and the nonspecifically captured off-target reads (i.e., from the non-coding region) (19, 35). Unfortunately, the information contained in the off-targets is too biased and incomplete. Our method utilizes the more complete SNP array data from the matched samples, which provides a more reliable and unbiased solution. iCNV uses a single hidden Markov model to jointly analyze data from all platforms. It assumes that those overlapping markers (i.e., exons and SNPs) share the same copy number and indeed use one platform to validate the calls from the other. In contrast, BMI-CNV systematically combines data sequences from multi-platforms and allows the overlapping markers to have different copy number states.

In this study, we developed a Bayesian statistical framework, which has several essential advantages over other modelling strategies. First, the nonparametric PSBP can relax the restrictive parametric assumption and allows flexible modelling of the complex high-throughput data. A common critique of the Bayesian method is its computational speed. In our framework, all conditional distributions implemented in the Gibbs sampling algorithm can be analytically derived, which guarantees efficient sample generation and fast computation. Second, the Bayesian framework enables great flexibility and possibility to incorporate prior relevant information such as the documented CNV hotspot information (20). Finally, the PSBP framework can be easily extended to accommodate the complex data dependence structure by replacing the independent weights with stochastic processes (e.g., Gaussian process) without sacrificing computational tractability (23).

Among the top associations with the TRICL study, the discovered significantly associated gene *LGALS9* was previously found to be a prognostic factor for lung cancer, low expression of which was correlated with poorer survival outcome (36). The significant amplification genes *HSPG2*, *EIF3E,* and *YTHDC2* also have been frequently described as oncogenes in diverse tumor types, including lung cancer, gastrointestinal cancers, and breast cancer (Li *et al.*, 2014; Chen *et al.*, 2018; Kalscheuer *et al.*, 2019; D. Wang *et al.*, 2019;). Although these genes were not significant after multiple comparison adjustments due to the small sample size, further large-scale studies are always needed to validate these potential findings.

Our method still presents limitations. First, it does not detect rare CNVs, as the power will be attenuated in the existence of a large proportion of non-carriers. Second, it will have low power to detect CNVs with similar proportions of duplications and deletions in the samples, which might be less likely to happen for germline variations. Our method may split those CNVs into several smaller deletions and duplications, as each CNV locus is equally likely to be assigned to deletion or duplication. Nevertheless, our method is innovative and opens a new door for integrative study to identify common CNVs, which has the potential to present a great impact on public health and basic biomedical science. Moreover, it can be extended to integrate other data types (e.g., gene expression data), which guarantees a future direction. In addition, our method may also be further extended to incorporate case-control status to directly identify disease-associated CNVs in a single model.

## Data availability

WES data from the 1000 Genomes Project are downloaded from the FTP site hosted at the EBI ftp://ftp.1000genomes.ebi.ac.uk/vol1/ftp/. SNP array data from the International HapMap project are available from the FTP site ftp://ftp.ncbi.nlm.nih.gov/hapmap/. The TRICL SNP array and WES data are available on request from the author CIA.

Supplementary Data are available at NAR online.

## Acknowledgements

We thank the reviewers in advance for their helpful and insightful suggestions and comments. We also acknowledge the support from the international TRICL consortium.

