## Supplementary text for "BMI-CNV: A Bayesian framework for multiple genotyping platforms detection of copy number variation"

Feifei Xiao

### Section A. Supplementary Methods

#### A.1 Data normalization and Standardization

We implemented a three-step median normalization procedure to mitigate the effect of three main observed sources of bias: exon length, GC-content and mappability (D'Aurizio *et al.*, 2016). Let $RC^{\mathrm{w}}_{\mathrm{k}'j}$ denote the raw read depth for exon $k'$ in sample $j$, each $RC^{\mathrm{w}}_{\mathrm{k}'j}$ was then normalized according to:

$$\widehat{RC}^{\mathrm{w}}_{\mathrm{k}'j} = RC^{\mathrm{w}}_{\mathrm{k}'j} \times \frac{m}{m_x} \tag{1}$$

where m is the overall median, $m_x$ is median of all the exons with the same values of exon length, mappability and GC-content. We applied this normalization procedure to both test and control samples. All pre-specified control samples were pooled together by averaging reads on each exon across all samples to form the common reference baseline. Finally, we calculated the $\log_2$-ratio of normalized read counts between test samples and the reference baseline (*log2R*).

To bring SNP array LRR and WES derived *log2R* to the same measuring scale, we standardized each via a robust scaling approach to produce $\hat{y}^{s}_{\mathrm{k}j}$ and $\hat{y}^{\mathrm{w}}_{\mathrm{k}'j}$. Specifically, let $y^{s}_{\mathrm{k}j}$ denote the LRR data corresponding to *k*-th SNP marker in sample *j*; let $y^{\mathrm{w}}_{\mathrm{k}'j}$ denote the normalized *log2R* for exon $k'$ in sample *j* (Rousseeuw and Croux, 1993), then

$$\hat{y}^{\mathrm{w}}_{\mathrm{k}'j} = \frac{y^{\mathrm{w}}_{\mathrm{k}'j} - median\left(y^{\mathrm{w}}_{\mathrm{k}'j}\right)}{\mathrm{interquartile}\left(y^{\mathrm{w}}_{\mathrm{k}'j}\right)} \; and \; \hat{y}^{s}_{\mathrm{k}j} = \frac{y^{s}_{\mathrm{k}j} - median\left(y^{s}_{\mathrm{k}j}\right)}{\mathrm{interquartile}\left(y^{s}_{\mathrm{k}j}\right)} \tag{2}$$

Where $\mathrm{interquartile}(.)$ equaled the difference between $75^{\mathrm{th}}$ and $25^{\mathrm{th}}$ percentiles.

##### A.2 Full model specification

We assumed a normal linear regression model, the $\phi_i$ was further modelled by a variable selection prior, where $G(.)$ is the PSBP.

$$Y_{ij} \sim N\left(Y_{ij}\middle|\phi_i\right) \tag{3}$$

$$\phi_i \sim \gamma_i \delta_0 + (1-\gamma_i)G(.); \delta_0 = (\mu_0, \tau_0) \tag{4}$$

$$G(.) = \sum_{l=1}^{L} \omega_l \, \delta_{\theta_l}(.); \theta_l = (\mu_l, \tau_l) \tag{5}$$

$$\omega_l = \Phi(\alpha_l) \prod_{r<l} \left(1 - \Phi(\alpha_l)\right); \alpha_l \sim N(\mu_\alpha, \tau_\alpha) \tag{6}$$

we introduced the latent variable $s_i$ and $z_{il}(s_i)$, $s_i = l$ denoted the $i$-th position was assigned to the $l$-th component,

$$s_i \sim multinomial(p_1, \dots, p_{L-1}) \tag{7}$$

$$z_{il}(s_i) = \begin{cases} N^-(\alpha_l(s_i), 1) \; l < s_i \\ N^+(\alpha_l(s_i), 1) \; l = s_i \end{cases} \tag{8}$$

For component specific parameters $\mu_l, \sigma_l$, we assumed conjugate normal and gamma hyperpriors:

$$\mu_l \sim N\left(\mu_\mu, \tau_\mu\right); \tau_l \sim Gamma(a_\tau, b_\tau) \tag{9}$$

For variable selection parameter $\gamma_i$, we assumed a Bernoulli-Beta conjugate prior:

$$\gamma_i \sim Ber(\kappa); \kappa \sim beta(a_0, b_0) \tag{10}$$

Assuming data were properly normalized and standardized, we adopted the following choices for the hyperparameters. For $G(.)$, $\mu_l = \{-5, -2, 0.1, 1, 2\}$; $\tau_l = \{0.4, 1, 1, 1, 1\}$; $\mu_\alpha = 0$; $\tau_\alpha = 1$; $\mu_\mu = 0$; $\tau_{mu} = 1$; $a_\tau = b_\tau = 0.5$. For variable selection prior, $a_0 = b_0 = 0.5$.

##### A.3 MCMC algorithm

Step 1. Update $s_i$ $for$ $i = 1, \ldots, m$, given $\gamma_i = 0$

$$Pr(S_i = l) = \frac{\omega_l N(y_{i.}|\mu_l, \tau_l)}{\prod_{l=1}^{L} \omega_l N(y_{i.}|\mu_l, \tau_l)} \tag{11}$$

where $\omega_l = \Phi(\alpha_l) \prod_{r<l} (1 - \Phi(\alpha_r))$.

Step 2. Update $z_{il}$ $for$ $i = 1, \ldots, m$ $and$ $l = 1, \ldots, L$

$$z_{il}(s_i) = \begin{cases} N^-(\alpha_l(s_i), 1) \; l < s_i \\ N^+(\alpha_l(s_i), 1) \; l = s_i \end{cases} \tag{12}$$

Step 3. Update $\alpha_l$ for $\mathrm{l} = 1, \ldots, \mathrm{L}$,

$$\alpha_l \sim N\left(\frac{\sum_{s_i>l} z_{il} + \mu_\alpha}{n_l + 1}, \frac{1}{n_l + 1}\right) \tag{13}$$

where $n_l = \sum_{i=1}^{m} 1(S_i \geq l)$.

Step 4. Update $\mu_l$ $for$ $l = 1, \ldots, L$,

$$\mu_l \sim N\left(\frac{\mu_\mu \tau_\mu + \tau_l \sum y(S_i = l)}{\tau_\mu + n_2 \tau_l}, \tau_\mu + n_2 \tau_l\right) \tag{14}$$

where $n_2$ is the number of elements in $y(S_i = l)$.

Step 5. Update $\tau_l$ $for$ $l = 1, \ldots, L$,

$$\tau_l \sim Gamma\left(a_\tau + n_2, b_\tau + 0.5 \sum (y - \mu_l)\right) \tag{15}$$

Step 6. Update $\gamma_i$ $for$ $i = 1, \ldots, m$

$$Pr(\gamma_i = 1) = \frac{a_i}{a_i + b_i} \tag{16}$$

$$a_i = \kappa \int \prod N(y(S_i = l)|\mu_l, \tau_l) f(\mu_l) \mathrm{f}(\tau_l) \, d\mu_l d\tau_l \tag{17}$$

$$b_i = (1 - \kappa) \prod N(y|\mu_0, \tau_0) \tag{18}$$

where $f(\mu_l)$ and $\mathrm{f}(\tau_l)$ are distributions of $\mu_l$ and $\tau_l$, respectively, the integration in equation (17) could be solved using the approximation:

$$p(y) = \frac{p(y|\mu_l, \tau_l) f(\mu_l) \mathrm{f}(\tau_l)}{p(\mu_l|y) p(\tau_l|y)} \tag{19}$$

Step 7. Update $\kappa$

$$\kappa \sim Beta(a_0 + \sum \gamma_i, b_0 + m - \sum \gamma_i) \tag{20}$$

##### A.4 SNP array and WES data processing

In the application of BMI-CNV to analyze 1000 genomes project and HapMap datasets (Altshuler *et al.*, 2010; Auton *et al.*, 2015). all Affymetrix raw CEL files were downloaded from website (ftp://ftp.ncbi.nlm.nih.gov/hapmap). We used the Affymetrix Power Tools and PennCNV package to generate LRR signals (Wang *et al.*, 2007). For WES data, the raw BAM files were downloaded from website (ftp://ftp.1000genomes.ebi.ac.uk). The BAM files were processed, sorted and filtered using SAMtools to generate raw read count (Li *et al.*, 2009). We then used the three-step normalization procedure described in Section A.1 to calculate the *log2R* data, 4 external samples: NA10851, NA18502, NA12272, NA19072 were selected as control samples.

In the application of BMI-CNV to analyze samples from international lung cancer study (TRICL) with both OncoArray data and WES data. (Amos et al., 2017). The OncoArray was designed from a list of 533,000 SNP markers. To retain high-quality genotype data, we applied the following quality control (QC) filters to remove (1) low quality samples (call rate<0.95); (2) unexpected duplicated and related samples (identical by descent (IBD)>0.2). Intensity data was obtained for each probe using GenomeStudio, genomic wave was adjusted by PennCNV. For WES data, QC procedures including base call quality recalibration variant filtering, genotypes refinement and Principal component analysis (PCA) of quality metric to exclude quality outliers. Kinship coefficient was also calculated to identify and exclude duplicated and related samples (Zheng et al., 2012). Raw read count data were then generated and normalized.

##### A.5 Post-calling CNV quality control (QC)

After obtaining the raw CNVs, to retain high quality CNV call, we first merged adjacent CNV calls (<5 markers). We then applied the following CNV pruning and filtering procedures including removing CNV calls that were (1) overlapped with centromeric regions; (2) < 3 exons or <10 SNPs, >100 markers and >500kb.

#### A.6 Numerical simulation

##### Data Simulation

For SNP array data, we simulated the log R ratio (LRR) and B allele frequency (BAF) from the normal distribution,

$$LRR \sim N(\mu_{LRR}, \sigma^2_{\mathrm{LRR}}) \quad (21)$$

$$BAF \sim N(\mu_{BAF}, \sigma^2_{\mathrm{BAF}}) \quad (22)$$

The empirical mean and variance values were provided by Illumina website (https://www.illumi na.com/documents) and summarized below (Table A.6.1).

For WES data, we spiked in the CNV signals by multiplying the raw read depth by a factor c/2, c was sampled from normal distribution,

$$c \sim N(\mu_{CNV}, \sigma^2_{CNV}) \quad (23)$$

Where the choices of mean and standard deviation (i.e. $\mu_{CNV}, \sigma_{CNV}$) parameters of different copy number states were summarized below (Table A.6.2).

##### Performance evaluation metrics

The performance of all the calling methods was assessed by precision rate, recall rate and F1 score measures. The precision rate which measured the proportion of CNV calls from methods that overlapped with the true CNV set was defined as, True positives/(True positives + False positives), while the recall rate which measured the proportion of true CNVs that were called by methods was defined as, True positives/(True positives + False negatives). The F1 score was defined as the harmonic mean of precision and recall rate which reflected the overall accuracy, $2\,\frac{precision*recall}{precision+recall}$.

**Table A.6.1. Copy number states and empirical parameter settings for SNP array data.**

| Copy Number State | LRR mean | LRR SD | BAF mean | BAF SD |
|---|---|---|---|---|
| Double Deletion | -5.00 | 2.00 | NA | NA |
| Single Deletion | -0.45 | 0.18 | 0, 1 | 0.03 |
| Normal | 0.00 | 0.18 | 0, 0.5, 1 | 0.03 |
| Single Duplication | 0.30 | 0.18 | 0, 1/3, 2/3, 1 | 0.03 |
| Double Duplication | 0.75 | 0.18 | 0, 1/4, 2/4, 3/4, 1 | 0.03 |

LRR: log R ratio; BAF: B allele frequency; SD: standard deviation.

**Table A.6.2. Copy number states and empirical parameter settings for WES data.**

| **Copy Number State** | $\mu_{CNV}$ | $\sigma_{CNV}$ |
|---|---|---|
| Double Deletion | 0.05 | 0.10 |
| Single Deletion | 1.00 | 0.10 |
| Single Duplication | 3.50 | 0.10 |
| Double Duplication | 6.00 | 0.10 |

### Section B. Supplementary Tables

#### Supplementary Table S1. Summary of BMI-CNV calling results on TRICL data.

| Methods | | N sample | N del | N dup | Del Mean size (markers) | Del Mean size (kb) | Dup Mean size (markers) | Dup Mean size (kb) |
|---|---|---|---|---|---|---|---|---|
| BMI-CNV | **Cases** | 1,075 | 66,705 | 62,440 | 12.91 | 32.30 | 8.38 | 33.21 |
| | **Control** | 917 | 68,560 | 55,478 | 14.00 | 30.64 | 9.09 | 31.72 |
| | **Total** | 1,992 | 135,265 | 117,918 | 13.46 | 31.46 | 8.72 | 32.51 |

N sample, number of samples; N del, number of deletions; N dup, number of duplications; Del, deletions, dup, duplications.

**Supplementary Table S2. Top significantly associated CNVs with lung cancer risk (P-value<0.005).**

| Gene | Stratum | CNV | Chr | Coordinate | Case:Cont | OR (95% CI) | P-value | P.adj |
|---|---|---|---|---|---|---|---|---|
| YTHDC2 | LUAD | Duplication | 5q22.2 | 112823818-112851198 | 37:22 | 2.88(1.62,5.12) | 0.0003 | 0.24 |
| MROH1 | LUAD | Duplication | 8q24.3 | 145246570-145509128 | 26:12 | 3.58(1.70,7.55) | 0.001 | 0.24 |
| NPEPPSP1 | LUAD | Deletion | 17q12 | 36351796-36413256 | 308:401 | 1.55(1.21,1.98) | 0.0005 | 0.07 |
| LOC101929950 | LUAD | Deletion | 17q12 | 36226155-36413730 | 308:401 | 1.55(1.21,1.98) | 0.0005 | 0.07 |
| HSPG2 | SQC | Duplication | 1p36.12 | 22156300-22170908 | 12:10 | 4.79(1.75,13.10) | 0.002 | 0.99 |
| EIF3E | SQC | Duplication | 8q23.1 | 109071781-109246041 | 33:54 | 2.19(1.31,3.64) | 0.003 | 0.99 |
| ACAD11 | SQC | Duplication | 3q22.1 | 132256840-132295990 | 20:27 | 2.69(1.39,5.20) | 0.003 | 0.99 |
| HMGA2 | SQC | Duplication | 12q14.3 | 66356901-66400689 | 15:11 | 3.50(1.44,8.53) | 0.005 | 0.99 |
| LGALS9 | SQC | Deletion | 17q11.2 | 25931519-26094865 | 11:17 | 4.14(1.65,10.38) | 0.002 | 0.21 |
| COG3 | SQC | Deletion | 13q14.13 | 46052970-46055553 | 19:30 | 2.74(1.39,5.38) | 0.004 | 0.21 |
| TESK2 | SQC | Deletion | 1p34.1 | 45836364-45864685 | 24:34 | 2.45(1.34,4.48) | 0.004 | 0.21 |
| FUBP1 | SQC | Deletion | 1p31.1 | 78413040-78415127 | 6:5 | 8.65(1.91,39.18) | 0.005 | 0.21 |
| TBC1D23 | SQC | Deletion | 3q12.1 | 99962992-100015266 | 12:4 | 5.64(1.66,19.15) | 0.005 | 0.21 |
| YTHDC2 | LC | Duplication | 5q22.2 | 112823818-112851198 | 54:22 | 2.43(1.42,4.18) | 0.001 | 0.28 |
| MROH1 | LC | Duplication | 8q24.3 | 145246570-145509128 | 35:12 | 2.89(1.41,5.93) | 0.004 | 0.56 |
| HSPG2 | LC | Duplication | 1p36.12 | 22156300-22170908 | 23:10 | 3.14(1.43,6.88) | 0.004 | 0.56 |
| SCARF2 | LC | Duplication | 22q11.21 | 20424584-20437826 | 56:21 | 2.19(1.27,3.77) | 0.005 | 0.56 |
| FAM230J | LC | Duplication | 22q11.21 | 18361721-18391105 | 56:21 | 2.19(1.27,3.77) | 0.005 | 0.56 |
| RIMBP3 | LC | Duplication | 22q11.21 | 18605815-18611919 | 56:21 | 2.19(1.27,3.77) | 0.005 | 0.56 |
| CRIPT | LC | Duplication | 2p21 | 46812338-46941301 | 26:8 | 3.28(1.43,7.49) | 0.005 | 0.56 |

P.adj is the adjusted P-values using Benjamini-Hochberg procedure; LUAD, lung adenocarcinoma; SQC, squamous cell lung cancer; LC, lung cancer; Chr, chromosome; Case, number of CNVs in cases; Cont, number of CNVs in controls; OR, odds ratio; 95% CI, 95% confidence interval.

### Section C. Supplementary Figures

**Supplementary Figure S1. Summary of length and frequency of BMI-CNV calling results on 1000 genome and HapMap project data.** Genomic length (in markers and kb) and population frequency of CNVs are compared between deletions and duplications. CNVs tend to be short in size with frequency less than 50%, whereas there is no difference between deletions and duplications.

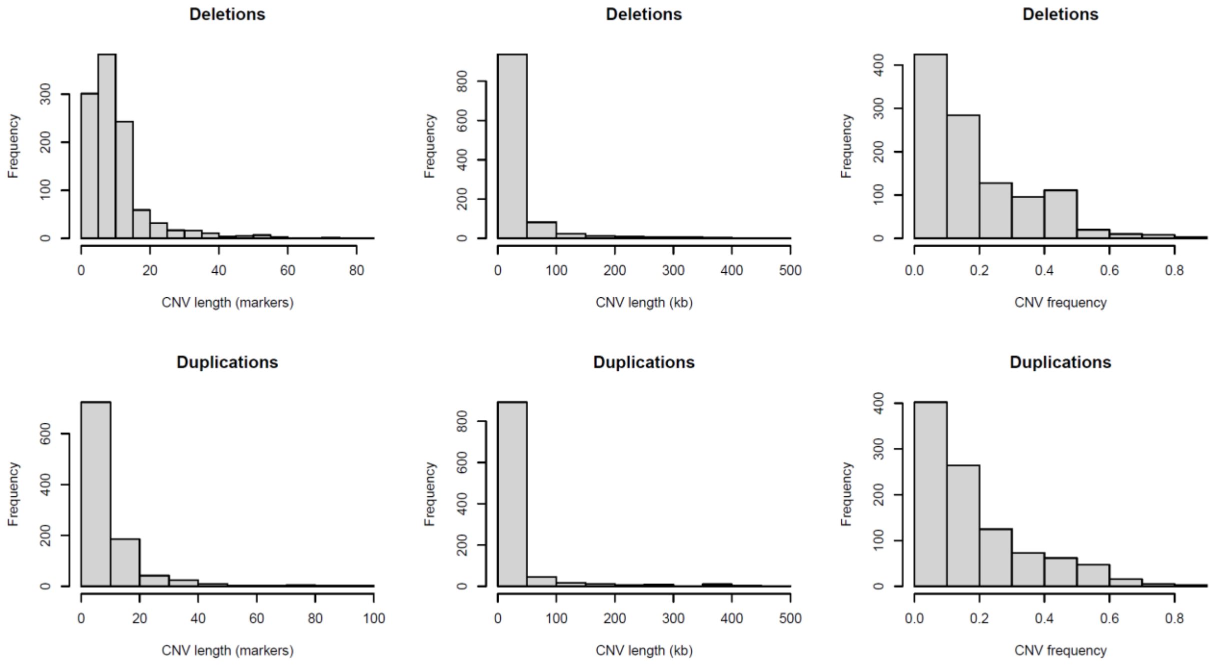

**Supplementary Figure S2. Case illustration of one common deletion region of chromosome 3.** The plot shows a common deletion region identified by BMI-CNV at position 52,404,727-52,425,389 of chromosome 3, this variant is shared across 27 samples. Each plot is a sample, x-axis is the genomic position and y-axis is the signal intensity (i.e. LRR or *log2R*). The mean signal intensities are shown by bold black lines. A clear shared deletion pattern is observed.

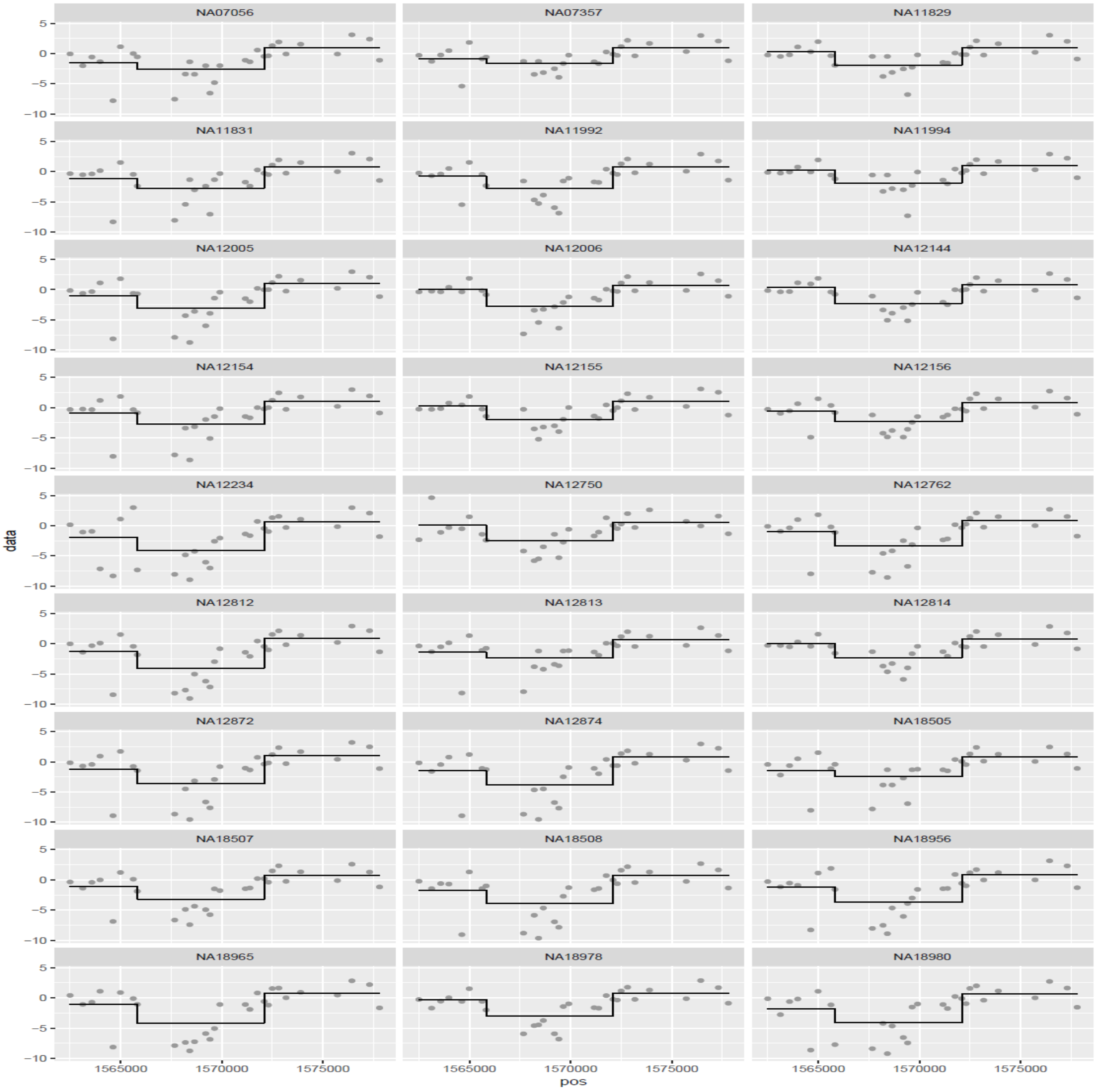

**Supplementary Figure S3. Comparison of length and frequency of BMI-CNV calling results on 1000 genome and HapMap project data under two different data integration strategies.** Genomic length (in markers and kb) and population frequency of CNV calling results under two data integration strategies are compared. Upper: data integration using exons and all SNPs; Lower: data integration using exons and intronic SNPs.

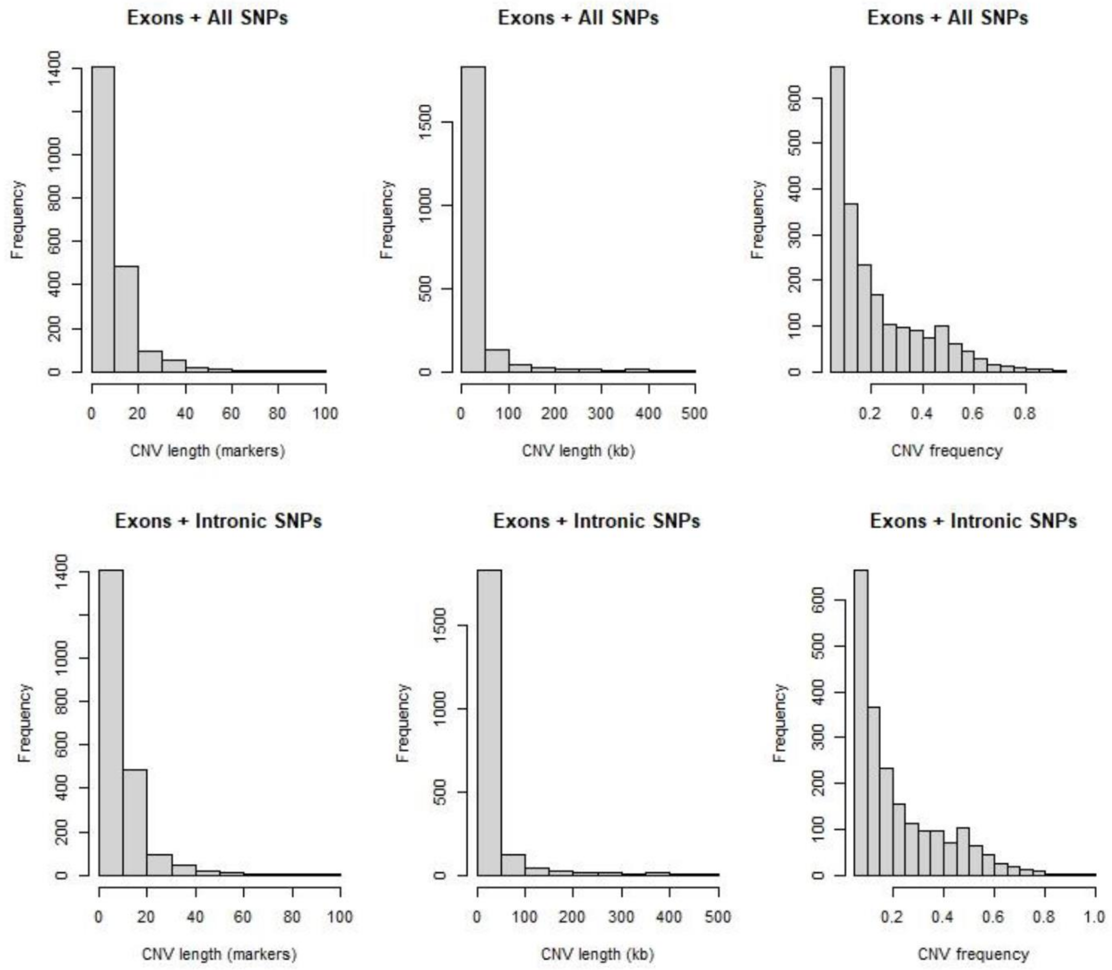

**Supplementary Figure S4. Data intensity of top lung cancer related CNVs.** The figure shows the data intensities of four lung cancer related CNV genes: (1) LGALS9; (2) HSPG2; (3) EIF3E; (4) YTHDC2. The mean signal intensity for each sample is shown by step bold line. For each gene, we included all carriers and other 50 randomly selected non-carriers. Vertical dashed lines depict regions identified by BMI-CNV, x-axis is the genomic position and y-axis is the signal intensity (i.e. LRR or *log2R*). All variants are valid CNV segments which show distinct data patterns from other non-carriers and adjacent regions.

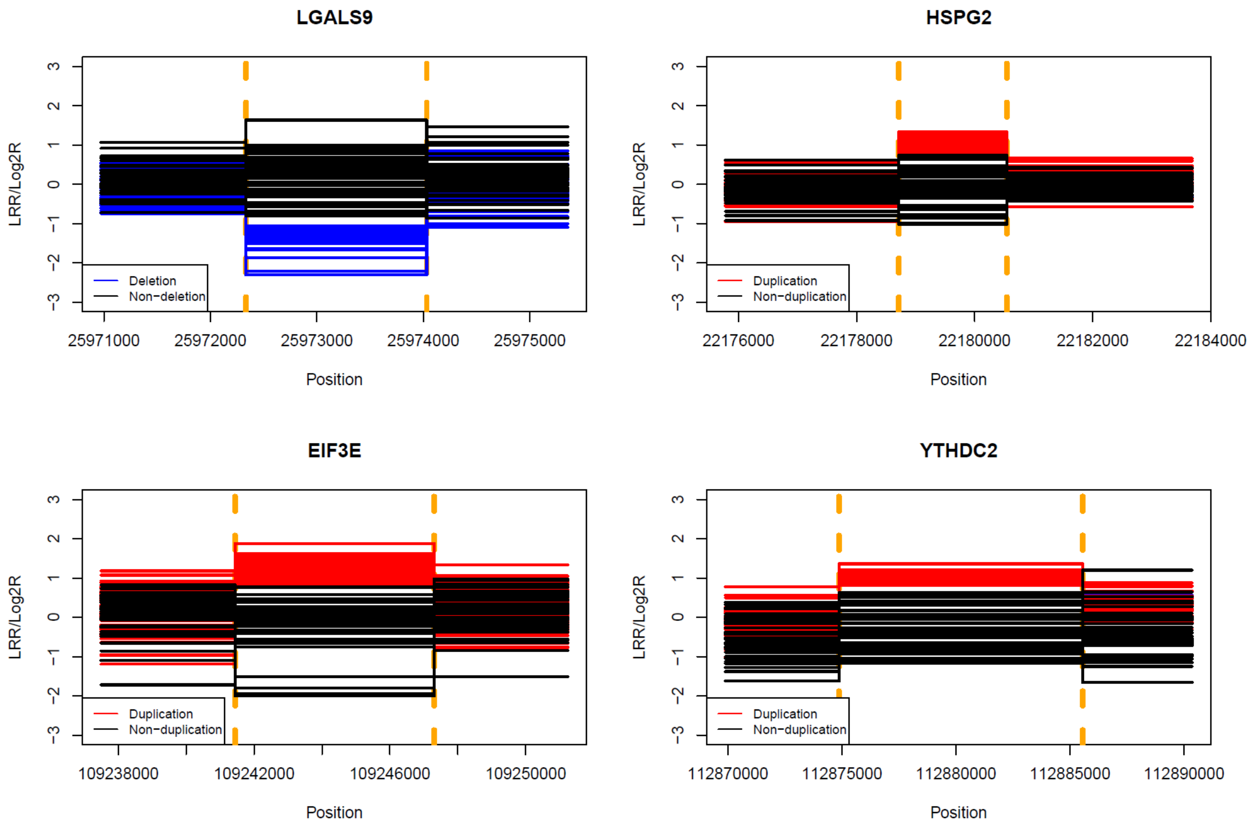
